# Temporal and Hemispheric Dynamics in Neural Processing of Auditory and Speech Stimuli Across Linguistic Complexity: A MEG Source Space Study

**DOI:** 10.1101/2024.12.05.626939

**Authors:** Patrick Krauss, Nikola Kölbl, Nadia Müller-Voggel, Stefan Rampp, Martin Kaltenhäuser, Konstantin Tzirdis, Achim Schilling

## Abstract

In this study we investigated the neural processing of auditory stimuli of varying complexity: a non-linguistic (pure tone), a simple linguistic (phoneme) and a complex linguistic (word) stimulus. We recorded brain activity of 30 healthy, righthanded participants using magnetoencephalography (MEG), and compared the resulting evoked fields (ERFs) in source space in three different time intervals, i.e. early (0-250ms), mid (250-500ms) and late (500-750ms) responses. Our results reveal a bilateral activation during early response and right-lateralized activation in the mid-phase for all stimuli. Hoewever, the late response exhibited lateralization variations. The pure tone predominantly activated the right hemisphere, consistent with pitch processing theories. The phoneme primarily engaged the left hemisphere, supporting its role in phonemic processing. Notably, the word elicited activation in both hemispheres, reflecting phonemic processing on the left and stress patterns on the right. These findings highlight the intricate interplay between temporal processing and hemispheric lateralization in speech perception, emphasizing the importance of stimulus complexity and temporal dynamics in understanding auditory and speech processing.

## Introduction

The brain’s capacity to process auditory stimuli, from basic tones to complex speech, showcases the intricate neural mechanisms involved in auditory perception and language processing. This ability is crucial for effective communication, enabling both sound recognition and interpretation within linguistic contexts. Through many functional MRI (Goucha & Friederici, 2015), but also (invasive) electroencephalography (EEG) (Metzger et al., 2020; Alday, 2019) and magnetoencephalography (MEG) (Tavabi, Obleser, Dobel, & Pantev, 2007; Hickok & Poeppel, 2007; Poeppel & Assaneo, 2020) studies, we know which brain regions are active in language processing, but the exact neuronal circuits and their temporal dynamics are still not fully understood. To unravel these neuronal processes, in recent years, neurolinguistic research has increasingly used natural continuous speech such as audio books as a stimulus (Schilling et al., 2021; Garibyan, Schilling, Boehm, Zankl, & Krauss, 2022; Koelbl, Schilling, & Krauss, 2023; Schüller, Schilling, Krauss, Rampp, & Reichenbach, 2023; Schüller, Schilling, Krauss, & Reichenbach, 2024). In this preliminary study, we want to decode human speech by analysing data from 30 healthy, righthanded subjects recorded with MEG, in an effort to apply the resulting knowledge to further studies of speech analysis. Given the high temporal resolution of MEG, we want to analyse not only localisation but also temporal dynamics of brain activity using event-related-fields (ERFs) in source space. For this purpose, we have chosen three auditory stimuli of different complexity: a nonlinguistic pure tone (5kHz), a simple linguistic phoneme (“i”), and a more complex linguistic stimulus, i.e. a single word (“gut”; German for “good”). In our analysis we find statistically significant differences in the spatio-temporal patterns evoked by the different stimuli in the primary auditory cortex, e.g. Heschl Gyrus (HG).

## Methods

Each stimulus was presented bilateral 30 times in a randomized sequence with a 1-second interval between onsets to 30 healthy, right-handed participants (15 females, average age: 22.9 ± 3) using a 248-channel MEG system (Magnes 3600WH, 4D-Neuroimaging). All measurements were approved by the Ethics Committee of the University Hospital Erlangen (No: 22-361-2). Data preparation involved using *MNE* software to identify and remove bad sensors, applying band-pass filters (1-20 Hz), down-sampling the data to 200 Hz and conducting independent component analysis to remove artifacts (Koelbl et al., 2023). Finally, the data were segmented into intervals from -0.5s to 1s corresponding to stimulus presentations. We calculated a vector volume source estimate (SE) for each subject and each stimulus utilizing a cortical source space (5mm grid) and the boundary element model of the template brain *fsaverage* from *FreeSurfer* (https://surfer.nmr.mgh.harvard.edu/) with method=“sLORETA”. We calculated the Grand Average for each stimulus over all subjects SEs, timeaveraged the resulting Grand Average SEs for three time intervals (early (0-250ms), mid (250-500ms), and late response (500-750ms)) and visualized them (Figure 1). Additionally, we examined activation differences in the primary auditory cortex, focusing on the HG, using label-time-courses within this region (Figure 2). For statistical evaluation we calculated paired Wilcoxon tests using the root-mean-square (RMS) amplitudes in the left and right HG (Table 1).

**Figure 1:**
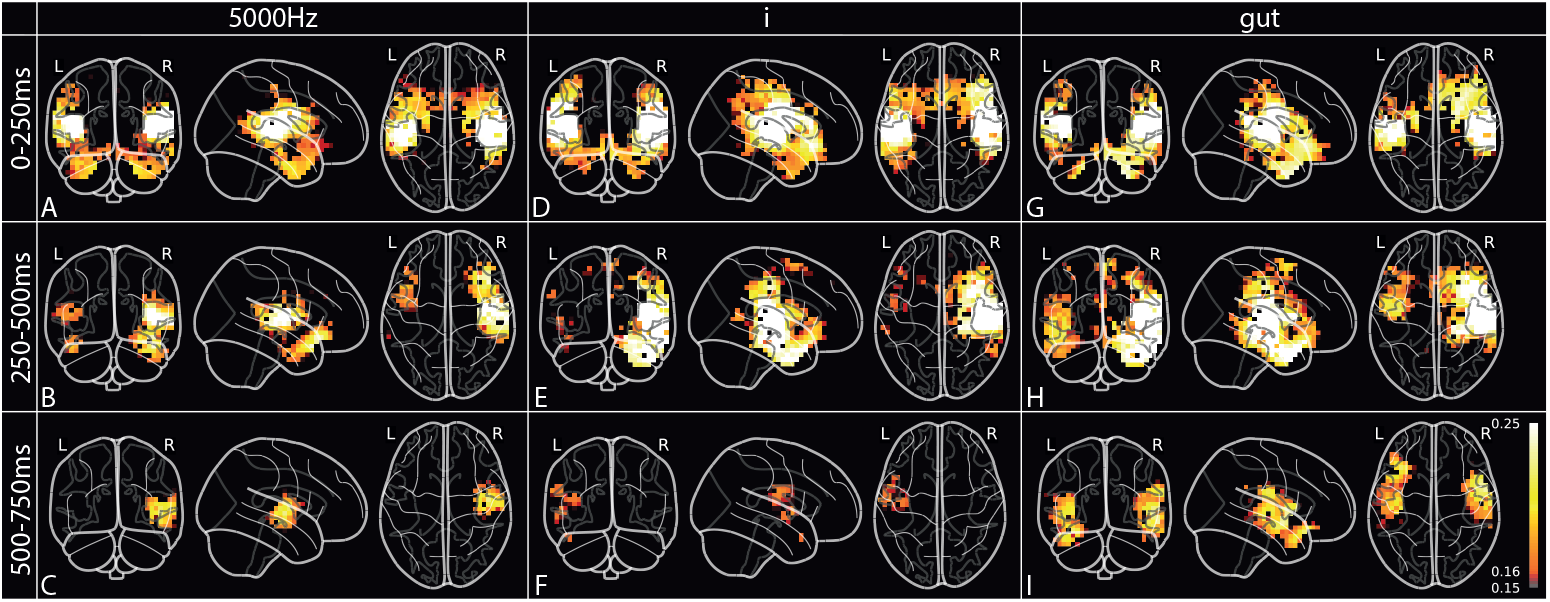
Grand Average Source Estimates over 30 participants for three stimuli (each presented 30 times) averaged over three time intervals. (A-C): Average brain response for pure tone 5kHz from 0-250ms, 250-500ms and 500-750ms. (D-F): Responses for phoneme “i”. (G-I): Responses for word “gut”.

**Figure 2:**
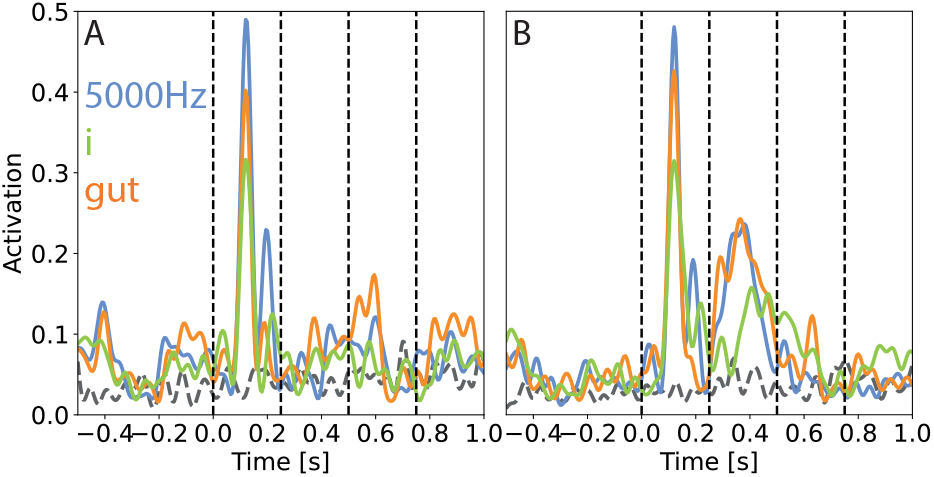
(A): Grand Average activity in left HG for three stimuli. (B): Same as (A), but in the right HG. Grey signal: Activity of random time-points.

**Table 1:** p-values of differences in activities in the left and right HG. Evaluated using a paired Wilcoxon Test with the RMS-amplitudes in three time intervals of three stimuli (5kHz, “i”, “gut”) of 30 participants. Bonferronicorrected significant p-values shown in red.

| left HG | 0-250ms | 250-500ms | 500-750ms |
| --- | --- | --- | --- |
| 5kHz vs. i | **0.0004** | 0.0155 | 0.0136 |
| 5kHz vs. gut | 0.0577 | **0.0016** | **0.0047** |
| i vs. gut | 0.0497 | 0.3387 | 0.0327 |
| right HG | 0-250ms | 250-500ms | 500-750ms |
| 5kHz vs. i | 0.0234 | **0.0050** | 0.9354 |
| 5kHz vs. gut | 0.3707 | **0.0001** | 0.0137 |
| i vs. gut | 0.0522 | 0.1981 | **0.0040** |

## Results

Our findings (Figure 1) indicate bilateral activation during the early response phase, despite amplitude differences possibly linked to audio volume. In the mid response phase, right-lateralized activation is prominent for all three stimuli, with larger amplitudes for the two linguistic stimuli. The late response phase unveiled distinct patterns: right-lateralized for the pure tone, left-lateralized for the phoneme and bilateral for the word, implying varied processing mechanisms. The average time courses in the left and right HG show similar patterns (Figure 2). The corresponding Wilcoxon tests of response amplitudes affirmed Bonferroni-corrected significant p-values (colored in red in Table 1), among others, between the non-linguistic and the complex linguistic stimuli in the mid time interval in both HGs and between the two linguistic stimuli in the late interval in the right HG.

## Discussion

Our research reveals the complexity of auditory and speech processing, showing specific activation patterns which differ particularly in the late processing phase depending on stimulus complexity. The predominance of right hemisphere activation could be related to the rhythmic presentation of stimuli, potentially reflecting the right hemisphere’s involvement in processing pitch and melody (Patterson, Uppenkamp, Johnsrude, & Griffiths, 2002). Conversely, left hemisphere activation for the simple linguistic stimulus (phoneme) during the late phase aligns with its role in phoneme processing (Desai, Liebenthal, Waldron, & Binder, 2008; Blumstein, Myers, & Rissman, 2005; Hyde, Peretz, & Zatorre, 2008) in particular in HG (Chait, Poeppel, De Cheveigné, & Simon, 2007). Bilateral activation for the more complex linguistic stimulus (word) underscores the intricate nature of language processing, requiring both interpretation of meaning and stress recognition. To deepen our understanding of speech perception’s neural underpinnings, future studies will have to explore additional brain areas. Additionally, machine learning could aid in identifying distinct activation clusters, offering deeper insights into the networks facilitating auditory processing and language comprehension.

## Acknowledgments

This work was funded by the Deutsche Forschungsgemeinschaft (DFG, German Research Foundation): grants KR 5148/2-1 (project number 436456810), KR 5148/3-1 (project number 510395418) and GRK 2839 (project number 468527017) to PK, grant TZ 100/2-1 (project number 510395418) to KT, and grant SCHI 1482/3-1 (project number 451810794) to AS.

